# GT-TS: Experimental design for maximizing cell type discovery in single-cell data

**DOI:** 10.1101/386540

**Authors:** Bianca Dumitrascu, Karen Feng, Barbara E Engelhardt

## Abstract

We present the Good-Toulmin like estimator via Thompson sampling, a computational method for iterative experimental design in multi-tissue single-cell RNA-seq (scRNA-seq) data. Given a budget and modeling cell type information across tissues, GT-TS estimates how many cells are required for sampling from each tissue with the goal of maximizing cell type discovery across samples from multiple iterations. In both real and simulated data, we demonstrate the advantages of GT-TS in data collection planning when compared to a random strategy in the absence of experimental design.

---

As both experimental and computational techniques advance, single-cell RNA-seq (scRNA-seq) allows for the characterization of cell type and cellular diversity at unprecedented high-throughout. Taking advantage of these developments, recent scientific efforts aim at the molecular profiling of all the cell types of an organism [Han et al., 2018, Regev et al., 2017]. These initiatives raise important questions regarding experimental design choices. First, given a budget, how many cells should one sample to maximize cell type discovery? Second, how should one prioritize among sampling organs or tissue types?

Both of these experimental design questions are related to the ecological problem of estimating the number of unseen species from a population. In the 1940s, the British naturalist Alexander Corbet spent two years collecting butterflies in Malaya, and he observed 118 butterfly species for which he had trapped only one specimen, 74 for which he had trapped two specimens, and 44 for which he had trapped three specimens. Following these trips, he wanted to estimate the expected number of distinct new species of butterflies he expected to find if he were to conduct a new expedition to Malaya.

Corbet posed this question to the famed mathematician Ronald Fisher who developed a parametric empirical Bayes method to estimate the expected number of new species with further sampling based on prior discoveries [Fisher et al., 1943]. Good further extended this work with the Good-Toulmin (GT) estimator [Good and Toulmin, 1956, Good, 1953]. Since this estimator was developed, many statistical approaches having been proposed, spanning parametric and nonparametric statistics and frequentist and Bayesian approaches [Orlitsky et al., 2016]. Variants of the problem exist in a variety of fields. In linguistics, the GT estimator has been applied to estimate the number of words in Shakespeare’s vocabulary [Efron and Thisted, 1976]; in genetics, it has been applied to estimate the number of unobserved genetic variants in the human genome [Ionita-Laza et al., 2009].

The history of the species discovery problem is rooted in a two-trial case over a single population of interest: *n* samples are collected from a population (first trial), and an expectation over the number of unseen species is estimated, if *c ∗ n* additional samples were collected, for a given extrapolation factor *c* (second trial). This setup has since been extended to an online learning setting over multiple populations [Battiston et al., 2016, Bubeck et al., 2013, Favaro et al., 2016, Good, 1953, Lijoi et al., 2007, Mao, 2004, Raghunathan et al., 2017]. These algorithms rely on solving the multi-armed bandit problem [Robbins, 1985], which balances exploration of the experimental choices with exploitation of current estimates of expected rewards – here, a new species. In the classical set-up, a gambler at a casino is presented with a row of slot machines (“one-armed bandits”) that each pay out a random reward according to their machine-specific probability distributions. The goal of the gambler is to maximize her winnings over a number of trials by deciding the sequence of machines to play; to do so, she must balance between exploiting the current estimates of expected arm rewards to select arms with the highest expected rewards and exploring undersampled arms in order to improve the estimates of the arms expected rewards.

In this work, we develop a principled approach to guiding the iterative selection of samples through single cell sequencing technologies when presented with the possibility of querying multiple tissues or sample sites. Motivated by multi-armed bandits, we consider this problem as a set of sequential trials where a scientist may choose a subset of tissue samples to assay, constrained by a budget. Each tissue or sample site represents an arm to be pulled, and upon choosing the tissue, the scientist can query a number of cells proportional to the maximum expected number of new cell type discoveries in the current experiments. The reward of each experimental trial is given by the number of new cell types uncovered in the chosen sample.

We propose the Good-Toulmin like estimator via Thompson sampling (GT-TS), a robust approach to experimental design for cell type discovery. Given a per trial budget, GT-TS leverages information across tissues to inform subsequent experiments in order to maximize cell type diversity and discovery. The GT-TS proceeds in two stages. First, a warm-up batch is selected from each possible tissue type, and an initial population-wise diversity metric is estimated using a Good-Toulmin like estimator (Methods, Fig. 1 a). The Good-Toulmin estimator quantifies the expected number of new cell type discoveries for each tissue type conditioned on the available samples from subsequent experiments. Second, given a budget *M* corresponding to the allowed number of samples to be collected, GT-TS iteratively and stochastically selects an experiment from a set of possible tissue types in proportion to their current Good-Toulmin estimate of expected rewards. We considered two variants of GT-TS, an *incidence case* and a *delayed abundance* case corresponding to different experimental design scenarios. The incidence case corresponds to a setting where, at each trial, all *M* cells are collected from one tissue type; in the delayed abundance case, the *M* cells are sampled from a subset of tissues, and the budget is partitioned across the tissues relative to the estimated probability of finding novel cell types in each of the tissues.

**Figure 1:**
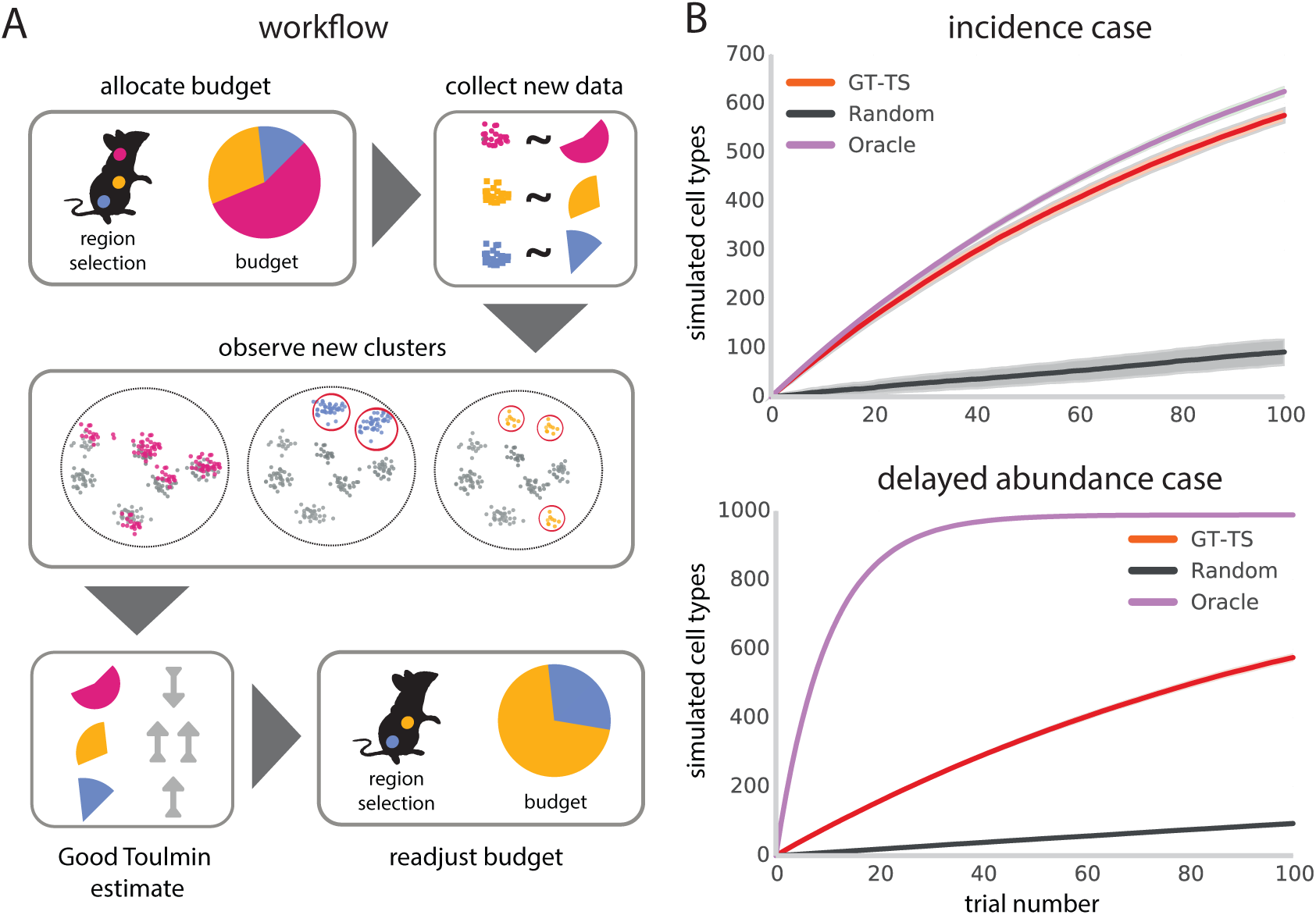
Panel A: Workflow for GT-TS. At each experiment, we select cells proportional to the Good-Toulmin estimate of new cell types; after each experiment, the estimates are recomputed to incorporate the current experiment. Panel B: Comparison of the average cell types discovered by the GT-TS, Random and Oracle algorithms on the heterogeneous simulated data set over 100 trials across 100 runs (standard deviation shown as shaded region). We consider the incidence case (top) and the delayed abundance case (bottom).

To evaluate the performance of GT-TS, we considered both simulated and existing single cell RNA-seq data. We compared our results to two relative strategies: a baseline Random strategy for which the probability of each tissue type being selected is equal, and an Oracle strategy, which makes the optimal arm choice. We demonstrated that GT-TS maximizes the number of cell types that can be discovered when compared with the Random strategy.

First, we generated a *needle-in-the-haystack* toy data set with 1, 000 cell types distributed across 10 tissues over 100 trials (Methods). The cell types were distributed across the tissue populations as follows: nine of the populations only contained species 1, while one population contained species one thousand unique species with uniform probability. We averaged the experiments over 100 runs. We found that the GT-TS algorithm outperforms the Random algorithm in both the Incidence case and the Delayed Abundance case (Fig. 1b). On this simulation, the improvement was dramatic, leading to up to six times more unique cell types discovered in the first 20 trials relative to the Random algorithm. In the Incidence case, GT-TS performs nearly as well as the Oracle strategy on these data.

We then applied GT-TS to a scRNA-seq data set from the Mouse Cell Atlas Project, a compendium of over 400, 000 cells from all of the major mouse organs, whose goal is to catalog murine cell types based on digital gene expression Han et al. [2018]. The large number of cells allowed us to simulate settings in which the cells were collected in smaller batches than in the actual experiments (Methods). We aggregated the 43 organs into 4 developmental stages: embryo, fetal, neonatal, and adult. The resulting data set is heterogeneous across age groups and organs, with most of the possible experiments representing organs or tissues in the adult population (Fig. 2a,b). When simulating iterative experiments, we sampled cells with replacement from each experimental category in the data set. GT-TS outperforms the Random strategy in both the Incidence case and the Delayed Abundance case (Fig. 2 b,c). In particular, GT-TS with Delayed Abundance identifies 20% more unique cell types after five trials as compared with the Random strategy (Fig. 2 d), approximately matching the arms selected by the Oracle algorithm (Figure Fig. 2 e).

**Figure 2:**
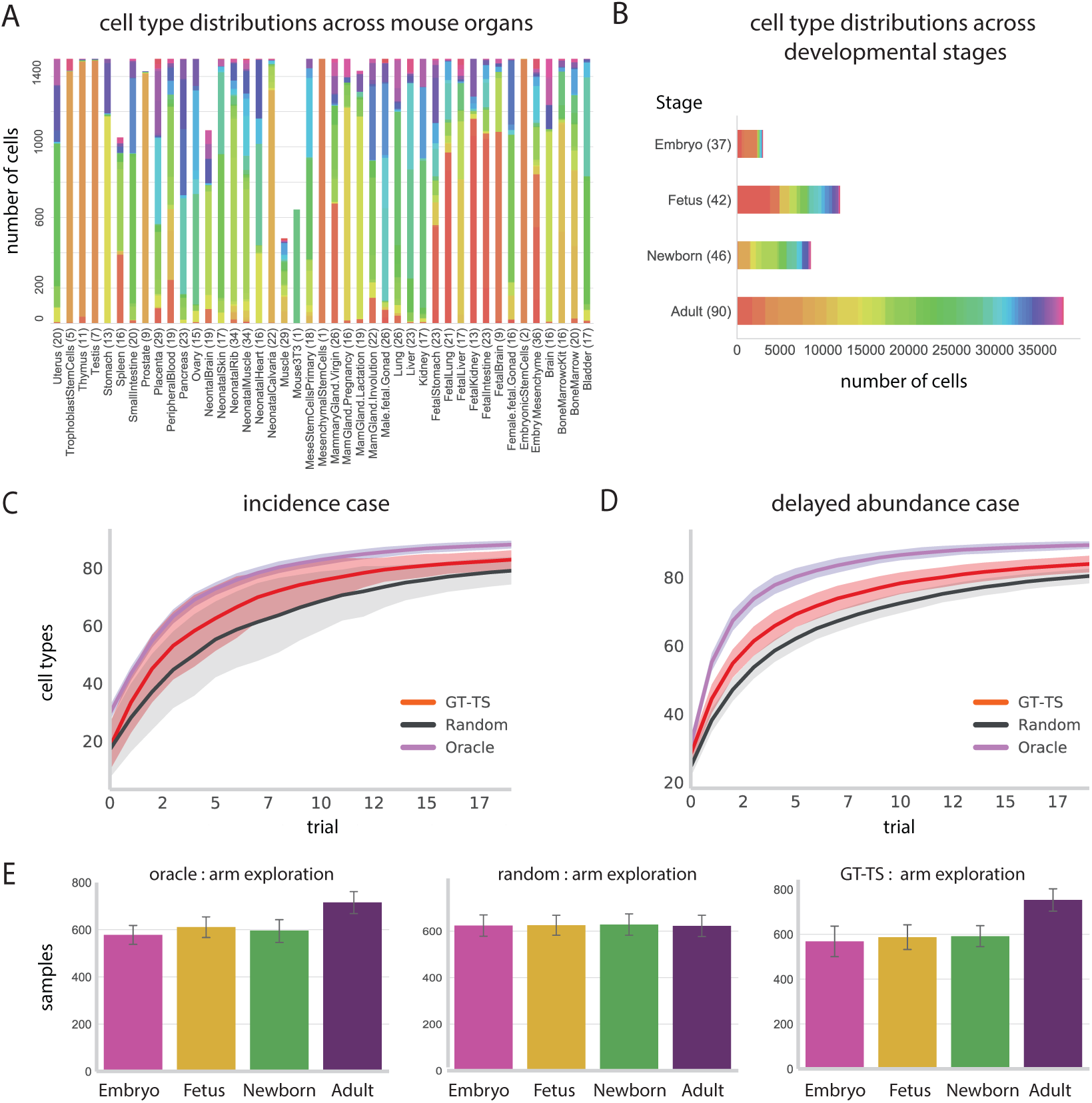
Panel A: Distribution of the cells found in 98 clusters across 43 organs in the original Mouse Cell Atlas data set. Each bar represents an organ, and each color represents a cluster. The *y*-axis indicates the number of clusters represented in each organ, and the *x*-axis represents the number of cells. Panel B: Distribution of the cells aggregated in 4 developmental stages. Each bar represents a stage, and each color represents a cluster. The *y*-axis indicates the number of clusters represented in each stage, and the *x*-axis represents the number of cells. Panel C: Comparison of the average clusters discovered by the GT-TS, Random, and Oracle algorithms on the Mouse Cell Atlas data set over 100 runs with 50 trials in the Incidence case. Panel D: Comparison of the average clusters discovered by the GT-TS, Random, and Oracle algorithms on the Mouse Cell Atlas data set over 100 runs with 50 trials in the Delayed Abundance case. Panel E: Comparison of the average and standard deviation samples drawn from each population by the GT-TS, Random,and Oracle algorithms on the Mouse Cell Atlas data set over 500 trials across 100 runs. GT-TS exhibits Oracle-like exploration behavior.

## Discussion

We develop an experimental design approach to iteratively select samples for single cell RNA-sequencing assays to maximize novel cell type discovery. Here, we assume that the cell type can be determined precisely using the scRNA-seq data; we know that this assumption is not true both for computational reasons—classification will be noisy—and for biological reasons–cell types are not discrete. Future work will incorporate richer models of cell type densities.

The difference in performance for both the Oracle and the sequential decision making strategies between the simulated and Mouse Atlas scRNA-seq data highlights an important point about population heterogeneity: GT-TS shows remarkable improvement over random sampling when the different arms or organs share cell types and have heterogeneous relative entropy in cell types. As heterogeneous cell types form tissues, and heterogeneous tissue types form organs, considering experiments on the level of organs may lead to more uncertainty in terms of cell type discovery than designing experiments on the level of tissue types. It is preferable then to specify the experiments at the most local level possible at the expense of increasing the number of experimental arms. Including additional information in the experiment such as spatial information will also increase the number of possible experiments and, consequently, design methods including GT-TS will be essential for systematic experiment prioritization. With this in mind, our approach is simple and can be immediately applied to improve the efficiency of experimental studies with alternative goals, such as designing sampling techniques for diversifying location-dependent tumor cell type heterogeneity Levitin et al. [2018]. We showed through empirical evaluation that our approach outperforms Random experiment selection when designing experiments with goal of maximizing cell type discovery.

## 1 Online Methods

### GT-TS workflow

GT-TS is a multi-armed bandit algorithm for experimental design that relies on two main concepts: i) an estimate of population’s cell type discovery potential based on a variant of the Good-Toulmin estimator, and ii) a classic Thompson Sampling (TS) routine [Abeille and Lazaric, 2017, Russo et al., 2017]. Pseudocode available in Supplementary Note 1.

### Good-Toulmin Estimates of tissue’s cell type discovery potential

Let Ω denote the set of tissues. Each of the *K* tissues (arms) corresponds to a probability distribution *D*_1_, …, *D_K_* over Ω, such that *D_j_*, *j* = 1,…,*K* represents the frequency of cell types in tissue *j*. At the initial stage, we have observed *n_j_* cells from the *j*th tissue, 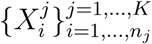. At a future experiment, we may choose to sample an additional *c_j_ ≤ n_j_* samples from the *j*th tissue, 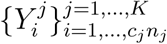, for an extrapolation factor *c_j_*. The number of new cell types discovered across all tissues is the statistic 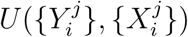, which is approximated as the sum of the number of new cell types discovered in each population *U_j_*(*{Y_i_}, {X_i_}*), where *j* = {1,…, *K*}

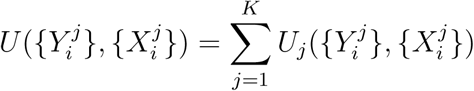

The truncated Good-Toulmin estimate Orlitsky et al. [2016] of *U_j_* is

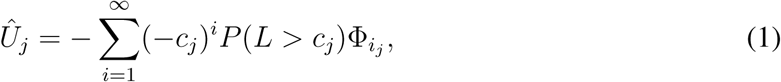

where 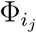 denotes the number of cell types observed in *i* cells in the sample from the *j*th tissue, and *L* is an independent random nonnegative integer.

### Experimental design via Thompson Sampling

Given a budget *M* constraining the number of cells that can be collected per trial and a set of *J* tissues to be explored, the algorithm iteratively selects a subset of tissues to sample from as well as the appropriate number of cells to sample. TS is a heuristic based on probability matching Thompson [1933]: the number of times an arm should be chosen should match its probability of being optimal. At each step, an arm is chosen based on its probability of being optimal according to the Bayesian posterior probability, which is then updated based on the given reward.

The probability that the *j*th tissue is chosen during a trial is based on the weight of its Good-Toulmin estimator 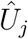. As the Good-Toulmin estimator can be negative and each arm must have a non-zero probability, we used a normalizing parameter *α* to set the probability of each arm as 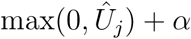; this results in the probability of the arm choice being

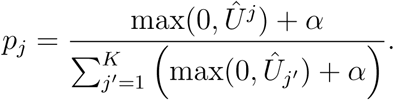

At each step, GT-TS allocates a budget *M* and calculates the desired number of samples from each population according to a probability mass function ***p*** = [*p*_1_,…,*p_K_*] for *c_j_* = 1 and a uniform random variable *L*. From the samples, *M* cell type labels are observed; the reward is the number of cell types that are novel discoveries. When labels are not available, a preferred clustering algorithm is employed and the cluster assignment become labels proxies that are iteratively updated.

### Hyperparameters

We set the hyperparameter *α* = 0.1 in both the simulated and real data. A larger *α* allows for more uniform exploration and higher variance in choosing the optimal tissues for sampling. The budget considered across experiments is *M* = 100. This budget is a low estimate of real life sequencing scenarios where collecting many samples of higher quality is expensive and experiments have to be prioritized, such as C1 and SMART-Seq technologies, which have throughput of 100-1000 cells Brennecke et al. [2013], Ziegenhain et al. [2017]. This budget is not higher because, in the Mouse Cell Atlas data, numbers greater than 100 often saturated the cell type diversity of a single tissue type and collapse the differences in performance between Oracle approaches and Random approaches. With larger sample sizes and greater cell type diversity, we hope to increase this budget in future work.

### Method Comparison

All results correspond to averaged results across 100 simulations or experiments.

### Mouse Cell Atlas Data

ScRNA-seq from Microwell-Seq in over 400, 000 cells from all of the major mouse organs representing > 800 cell types [Han et al., 2018]. Here, we define cell types based on digital gene expression. The Mouse Cell Atlas selected 60, 000 high-quality, batch corrected cells sampled from the complete data set that include 43 tissues and 98 major clusters of cell types. Note that this removed 87.5% of the diversity of cell types, and rare cell types in particular, limiting the complexity of the underlying data. Some cell type clusters presented significant multi-tissue contributions; for example, liver and muscle include cells from the hematopoietic cell cluster. Differential gene expression was performed by unsupervised clustering with the scRNA-seq data analysis tool Seurat [Butler et al., 2018].

## Competing Interests

The authors declare that they have no competing financial interests.

